# An Activator Locks Kv7.1 Channels Open by Electro-Mechanical Uncoupling and Allosterically Modulates its Pore

**DOI:** 10.1101/2021.02.11.430741

**Authors:** Melina Möller, Julian A. Schreiber, Mark Zaydman, Zachary Beller, Sebastian Becker, Nadine Ritter, Eva Wrobel, Nathalie Strutz-Seebohm, Niels Decher, Jianmin Cui, Nicole Schmitt, Martina Düfer, Bernhard Wünsch, Guiscard Seebohm

## Abstract

Loss-of-function mutations in K_v_7.1 often lead to long QT syndrome (LQTS), a cardiac repolarization disorder associated with increased risk of arrhythmia and subsequent sudden cardiac death. The discovery of agonistic *I_Ks_* modulators may offer a new potential strategy in pharmacological treatment of this disorder. The benzodiazepine (*R*)-L3 potently activates K_v_7.1 channels and shortens action potential duration, thus may represent a starting point for drug development. However, the molecular mechanisms underlying modulation by (*R*)-L3 are still unknown. By combining alanine scanning mutagenesis, non-canonical amino acid incorporation, voltage-clamp electrophysiology and fluorometry, and *in silico* protein modelling, we showed that (*R*)-L3 not only stimulates currents by allosteric modulation of the pore domain but also alters the kinetics independently from the pore domain effects. We identified novel (*R*)-L3-interacting key residues in the lower S4-segment of K_v_7.1 and observed an uncoupling of the outer S4 segment with the inner S5, S6 and selectivity filter segments. Summarizing, we provide structural and functional evidence for two independent K_v_7.1 activating mechanisms by a single modulator.

## Introduction

K_v_7.1 channels are the founding members of voltage-gated K_v_7 delayed rectifier potassium channels (K_v_), that play important functions in various tissues including epithelia, brain, heart and inner ear organs ^1, 2^. In native tissues, K_v_7.1 α-subunits (encoded by *KCNQ1*) can interact with various ancillary subunits or modulatory proteins, which constitute the biophysical properties of the channel producing functionally distinct K^+^-currents ^3–7^. In cardiac myocytes, co-assembling of the pore-forming α-subunit and its regulatory β-subunit KCNE1 (mink, IsK) generates the slow component of the cardiac delayed rectifier potassium current *I*_Ks_ that is critical for the repolarization of the cardiac action potential ^8, 9^. Loss-of-function mutations in K_v_7.1 can lead to prolonged cardiac repolarization and cause long QT syndrome 1 (LQTS1), a genetically heterogeneous cardiac arrhythmia that is characterized by a prolonged ventricular repolarization phase ^10–12^. Recently, diabetes was linked to *KCNQ1* as well ^13–17^. Furthermore, K_v_7.1 channels are important in many other organs and contribute to different physiological functions ^1, 2, 4^. Thus, K_v_7.1 channel modulators hold the potential for development of new pharmacological strategies in the treatment of many diseases like cardiac arrhythmias, diabetes, diarrhea, impaired function of thyroid gland and others.

Homomeric K_v_7.1 channels exhibit activation upon membrane depolarization and undergo delayed partial inactivation. This delayed slow inactivation cannot be observed directly upon channel activation but becomes observable when the membrane is stepped to hyperpolarized potentials after extended channel (in)activation and channels switch from inactivated to activated states before closing. A characteristic hook in tail can thus be seen in the current traces. These characteristics of Kv7.1 gating have been proposed to be the result of at least two open states that need to be occupied during gating process, from which the channel can enter flicker states with several subconductive states ^18–20^.

The prototypic high potency activator of K_v_7.1 is the benzodiazepine (*R*)-L3 (L-364,373, Figure 1a), that leads to increased *I*_Ks_ current amplitude, shortened action potential duration in guinea pig cardiac myocytes ^21^ and suppresses early after-depolarizations in rabbit ventricular myocytes ^22^. The K_v_7.1 channel modulation by (*R*)-L3 is associated with pronounced alterations of gating parameters ^23, 24^. (*R*)-L3 appears to arrest the channel in closed and open states and therefore prevent the channel from entering the inactivated state, which is appreciable as an absent hook in the tail currents ^23^. Thus, the presence of (*R*)-L3 seems to reduce or eliminate a particular gating transition towards subconductive states. The β-subunit KCNE1 increases channel conductance, slows channel activation and deactivation and markedly reduces macroscopic inactivation as well ^8, 9, 18, 25–28^. Thus, (*R*)-L3 and KCNE1 cause somewhat similar effects on K_v_7.1 channel. However, in K_v_7.1/KCNE1 heteromeric channels the hook in the tail can be made visible by replacing the conducted ion K^+^ by Rb^+ 18^.

**Figure 1.**
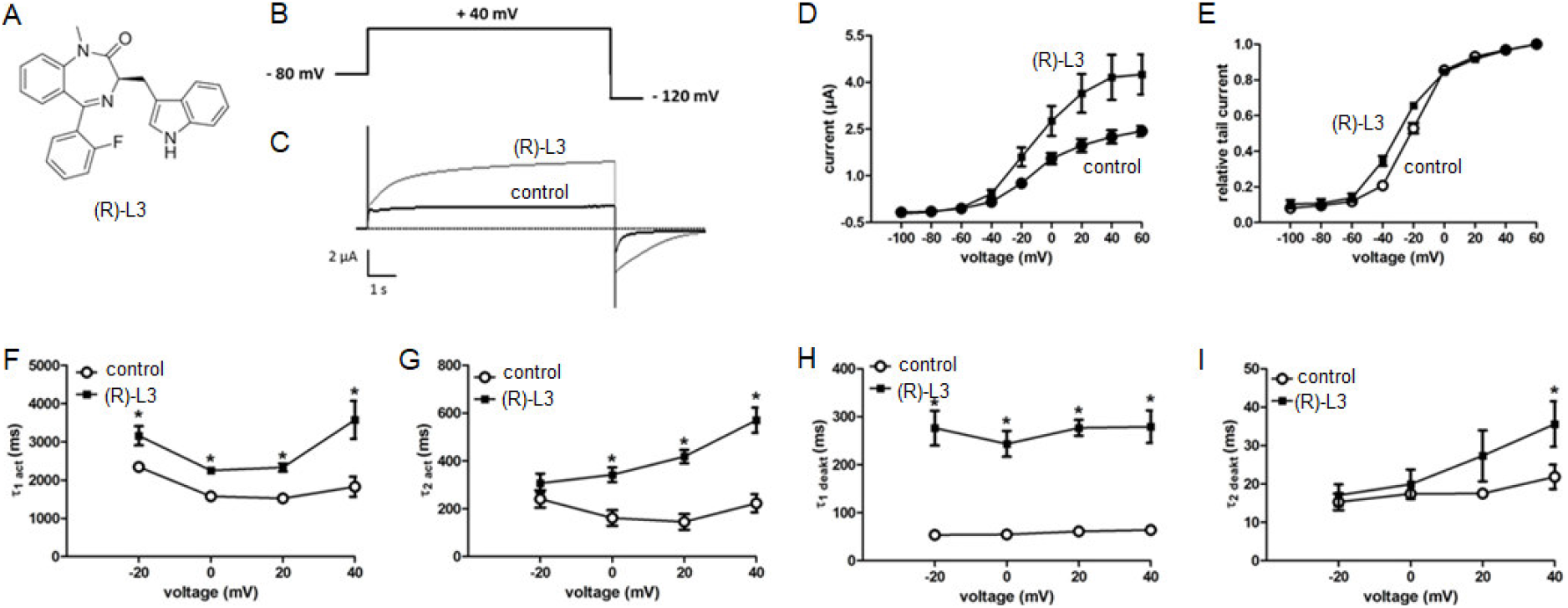
(*R*)-L3 activates and slows the rates of activation and deactivation of cloned hK_v_7.1 channels expressed in *Xenopus* oocytes. (A) Chemical structure of (*R*)-L3 (B) Pulse sequences for voltage-clamp experiments. (C) Effect of (*R*)-L3 on K_v_7.1 currents, recorded in an oocyte by a 7-second pulse to potentials of −100 mV to +60 mV from a holding potential of −80 mV. Currents were recorded in control solution containing 0.1 % DMSO followed by perfusion with 1 μM (*R*)-L3 containing solution. (D) Current-voltage relationship for K_v_7.1 current in the absence and presence of 1 μM (*R*)-L3. (E) Voltage dependence of current activation determined from peak tail currents measured at −120 mV. Currents were normalized to the peak tail currents elicited after a pulse to +40 mV (n=15 ± SEM). (F-I) Kinetics were evaluated at +40 mV in both absence and presence of 1 μM (*R*)-L3 and fitted by two exponential function. Time constants for fast and for slow component (G) of K_v_7.1 activation. (H) Time constants for fast and for slow component (I) of K_v_7.1 deactivation (n=15 ± SEM; *p<0.05).

Given the therapeutic potential of (*R*)-L3-like compounds in LQTS, there is a strong interest in defining the compound binding site and in understanding the molecular mechanism of action. Previous data revealed that specific residues in the K_v_7.1 pore domain interact with (*R*)-L3 allowing the compound to slow K_v_7.1 channel kinetics and increase the ion channel current ^23^. However, the precise molecular mechanism of channel modulation remained elusive. Interestingly, it was shown that the K_v_7.2 channel opener NH29 seems to bind in the core of the voltage-sensing domain and thereby operating via an interaction with the voltage-sensor. Thus, this study suggests that the voltage-sensor domain (VSD) might be an important structural target to develop specific Kv7 channel agonists ^29^. We hypothesized that since (*R*)-L3 changes the activation and deactivation kinetics and the voltage-dependence of K_v_7.1 activation, (*R*)-L3 may also physically interact with the VSD.

To address this question, we systematically probed the lower S4 segment of K_v_7.1 via site-directed mutagenesis and subsequently analyzed (*R*)-L3 effects on the mutated channels. To gain insights in the mechanism of action, we performed experimental data guided 3D-modeling of K_v_7.1 and (*R*)-L3 to identify the complete (*R*)-L3 binding site. Subsequent molecular dynamics simulations were utilized to assess the stability of drug binding and to explore the binding mode of the drug.

## Results

In this study, we extend our analysis of the putative (*R*)-L3 binding site towards the VSD by two-electrode voltage-clamp experiments using hK_v_7.1 expressing *Xenopus laevis* oocytes. In agreement with our previous electrophysiological studies, we find a robust K_v_7.1 agonism by (*R*)-L3 ^23^. The effects of (*R*)-L3 on K_v_7.1 current amplitude and kinetics are summarized in Figure 1. Specifically, the current amplitude was significantly increased and the activation as well as the deactivation were significantly slowed (Figure 1D-1I). In addition, the voltage-dependence of activation was mildly but not significantly shifted to more negative voltages as described before (Figure 1D, 1E) ^23^.

We tested whether changing the permeating ion can restore the hook in the tail also under presence of (*R*)-L3. High extracellular K^+^ or Rb^+^ (100 mM) partially restored the peak amplitude of the K_v_7.1 tail current (Figure 2A-B). However, as the extent of channel tail current amplitude activation heavily depends on the permeating ion, the (*R*)-L3 tail current amplitude stimulation is much reduced in high K^+^ and Rb^+^ compared to that in ND96 extracellular solution (Figure 2C). The mildly but not significant shift in voltage-dependence of activation as well the slowing of activation and deactivation by (*R*)-L3 is maintained in high K^+^ and Rb^+^ solution (Figure 2D-I).

**Figure 2.**
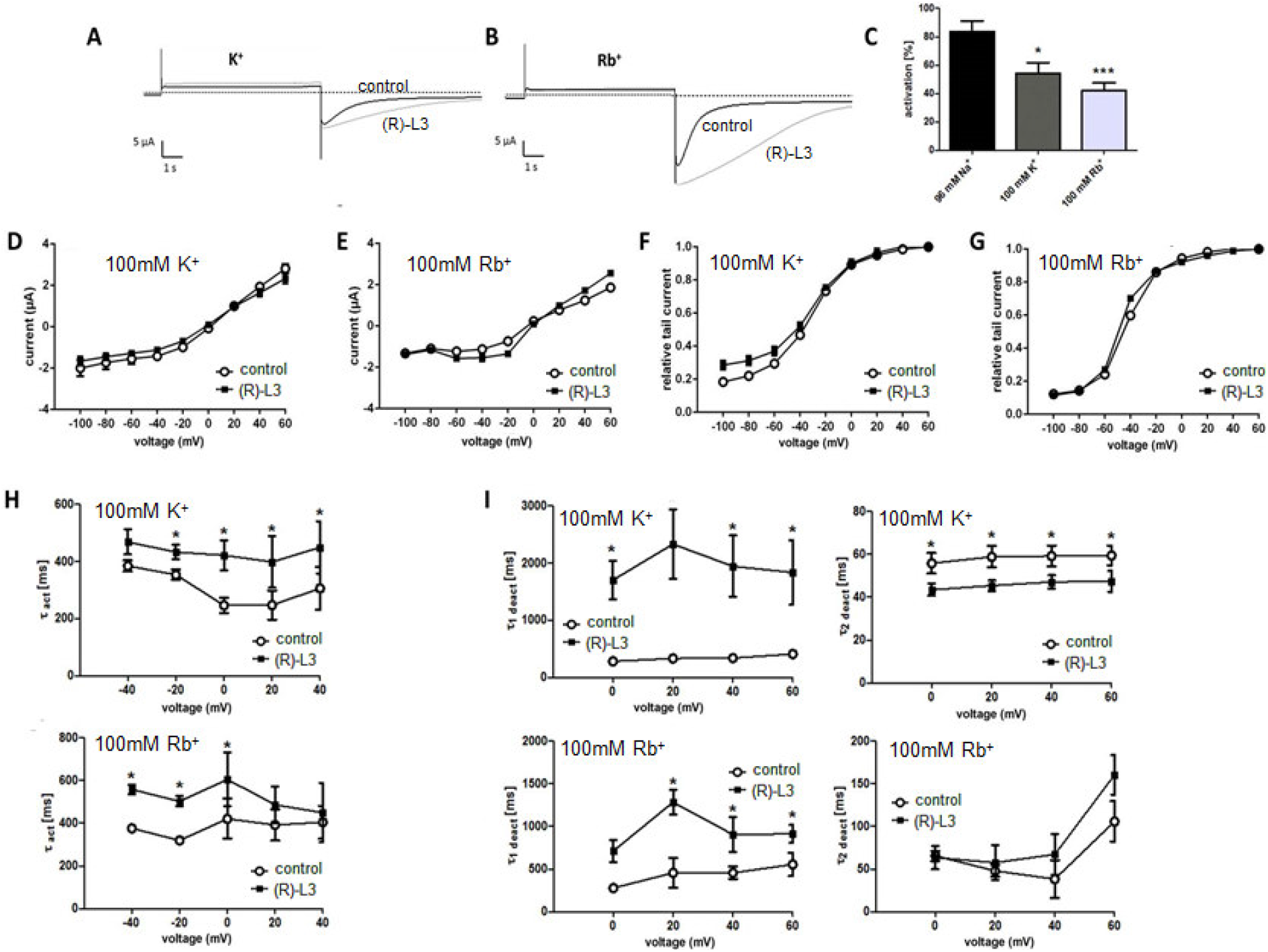
Effect of (*R*)-L3 on K_v_7.1 in high potassium and rubidium (100 mM each). (A-B) Representative current traces in high potassium (A) and rubidium (B) before and after addition of 1 μM (*R*)-L3. Activation of K_v_7.1 channel currents by (*R*)-L3 expressed as percent change in current in high sodium, potassium and rubidium. To illustrate the extent of activation by (*R*)-L3, the peak current activated by a 7-sec pulse to +40 mV was scaled to match the peak current recorded after the addition of 1 μM (*R*)-L3 (n=15 to 18 ± SEM). (C) Activation of K_v_7.1 channel currents by 1 μM (*R*)-L3 expressed as percent change in current in high sodium, potassium and rubidium (n=15 to 18 ± SEM). Current-voltage relationship for K_v_7.1 current before and after addition of 1 μM (*R*)-L3 in high potassium (D) and in high rubidium (E). Voltage dependence of current activation determined from peak tail currents measured at −120 mV in high potassium (F) and rubidium (G). Currents were normalized to the peak tail currents elicited after a pulse to +40 mV (n=15 ± SEM). Kinetics were evaluated at +40 mV in both absence and presence of 1 μM (*R*)-L3 and fitted by one and two exponential functions. (H) Single time constant of K_v_7.1 channel activation in high K+ and high Rb+ (n=13 ± SEM; *p<0.05). (I) Time constants for fast and slow component of K_v_7.1 channel deactivation in high K+ and high Rb+ (n=13 ± SEM; *p<0.05). The closed squares mark control data and open circles indicates (*R*)-L3 results.

In order to analyze if the (*R*)-L3 interaction affects the voltage-dependent movements of the VSD, we performed voltage clamp fluorometry experiments. We utilized a K_v_7.1-construct in which two extracellularly accessible cysteines (C214A/C331A) were removed and a novel cysteine (G219C) was introduced in the VSD for site-specific attachment of Alexa 488-C5 maleimide. This approach allows for monitoring of voltage-dependent VSD-movements ^30, 31^. As illustrated in Figure 3, we found that the fluorescence voltage (FV) curve, reflecting the voltage-dependend activation of VSD, is shifted toward more negative potentials by (*R*)-L3, indicating that (*R*)-L3 interactions modulate the VSD movement. Recently, it was shown that phosphoinositides (PI(4,5)P_2_) are required to couple VSD-activation to the conformation of the pore ^31^. We tested if the effects of (*R*)-L3 on VSD movement are likely direct or indirect by repeating the voltage-coupled fluorescence (VCF) experiment in cells expressing a voltage-sensitive lipid phosphatase (CiVSP), which depletes membrane PIP_2_ when activated ^32^. Any effects of (*R*)-L3 on the pore conformation, which may affect the VSD movement indirectly through the VSD-pore interactions, should be largely reduced or completely eliminated by such PIP_2_ depletion ^31^. We found that (*R*)-L3 shifts the FV curve similarly in the presence and absence of endogenous PIP_2_ (Figure 3E) suggesting that the effects of (*R*)-L3 on VSD activation are through a direct interaction with the VSD.

**Figure 3.**
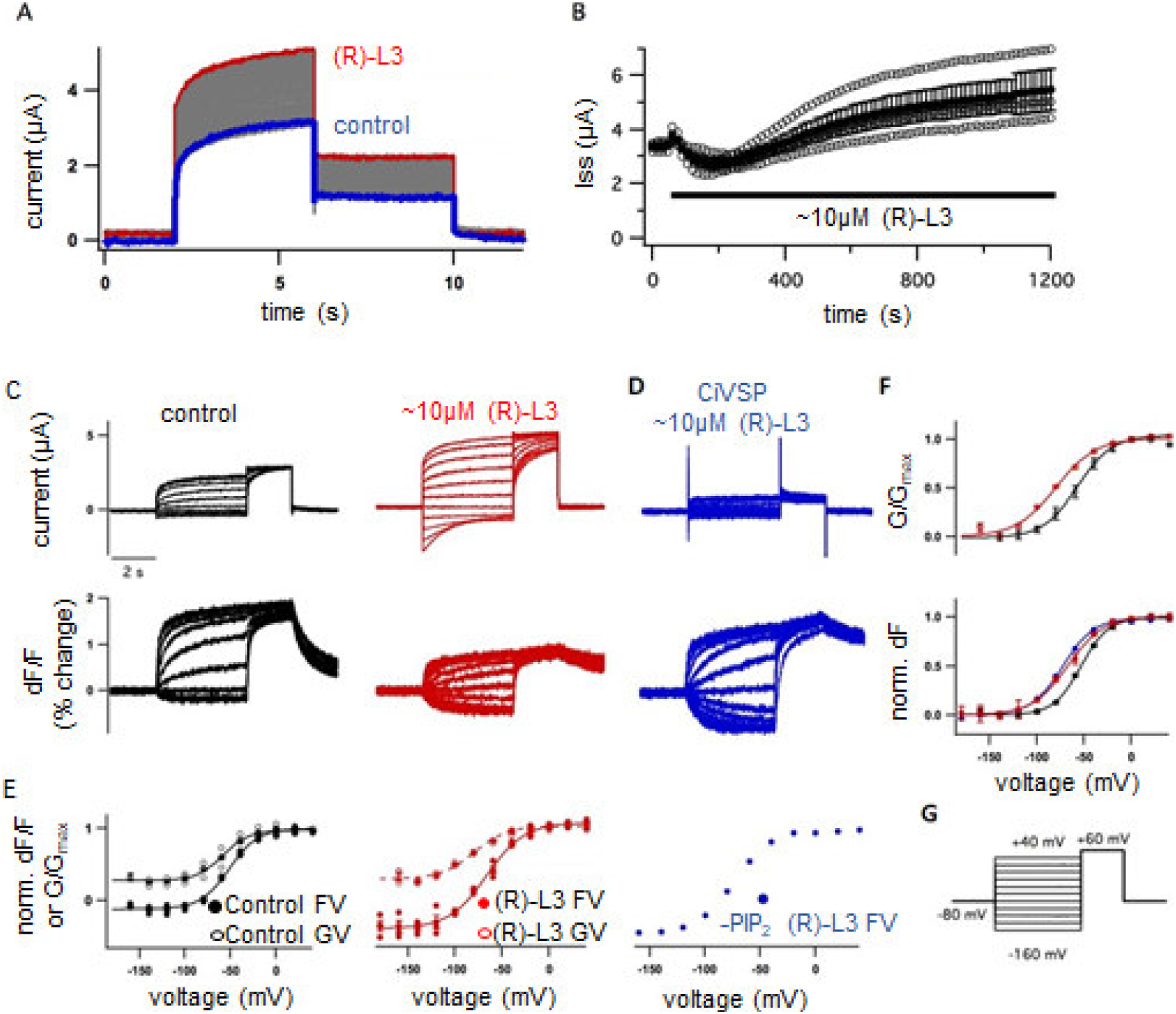
(*R*)L-3 potentiates psWT (C214A/G219C/C331A) Kv7.1 current and left shifts the voltage-dependence of VSD activation in the presence of endogenous PIP_2_ and following PIP_2_ depletion by CiVSP. (A) Whole cell currents from an oocyte expressing psWT Kv7.1 labeled with Alexa 488 C5 maleimide. Every 15 seconds, the membrane voltage was pulsed from the. −80 mV resting potential to +60 mV for 4 seconds, followed by 2 second tails at −40 mV. Currents before (blue) and after (red) a bolus of (*R*)-L3 was added to the bath (final concentration ~10 μM). (B) Steady-state current at +60 mV versus time during application of (*R*)-L3 (indicated by bar). Oocytes expressing psWT Kv7.1 alone (black, red) or with CiVSP (blue) were labeled with Alexa 488 C5 maleimide. (C) Whole oocyte current (top) and relative fluorescent signal changes (bottom) from a single oocyte before (black) and after (red) exposure to ~10 μM (*R*)-L3. (D) Whole oocyte current (top) and relative fluorescent signal changes (bottom) from a single oocyte expressing CiVSP following exposure to ~10 μM (*R*)-L3 and PIP_2_ depletion by a train of high voltage pulses. (E) Normalized fluorescence-voltage (FV) and conductance-voltage (GV) curves of individual and averaged data records. (F) Offset and normalized GV (top) and FV (bottom) curves. (G) Voltage protocol (sweep interval = 15 s).

In our previous study, we mapped (*R*)-L3 binding to the interface between the pore domain α-helices S5 and S6 and the outer face of S5 ^23^. Indeed, the large benzodiazepinic region of (*R*)-L3 was proposed to attach to the outer S5, that in turn is expected to interact with the voltage sensor domain of K_v_ channels in open or inactivated states ^33^. Thus, the (*R*)-L3 binding site may overlap with the interacting surface of the VSD and the pore domain. Specifically, the lower S4 helical transmembrane segment and the benzodiazepinic part may interact with similar regions of the lower S5. In order to test this hypothesis, we performed alanine-scanning mutagenesis of the lower S4 transmembrane segment (residues 235-241) and test for effects on modulator activity on mutant channels (Figure 4). We found that (*R*)-L3 effects on current amplitude could be non-significantly altered (I235A, M238A, L239A and V241A), increased (R237A and H240A) or decreased (L236A) by the respective amino acid exchange (Figure 4B). As the modulator slows activation and deactivation kinetics in wt channels the effect on kinetics of mutant channels was assessed. The effect of the modulator on activation was largely reduced or abolished in all mutant channels except H240A. However, slow deactivation was only unaffected by (*R*)-L3 in M238A (Figure 4B). Thus, only substitution of methionine 238 by alanine eliminates all different (*R*)-L3 effects on gating kinetics. Interestingly, the previous coupled effects on channel kinetic and current amplitude by (*R*)-L3 become more uncoupled by the examined mutations of the lower S4.

**Figure 4.**
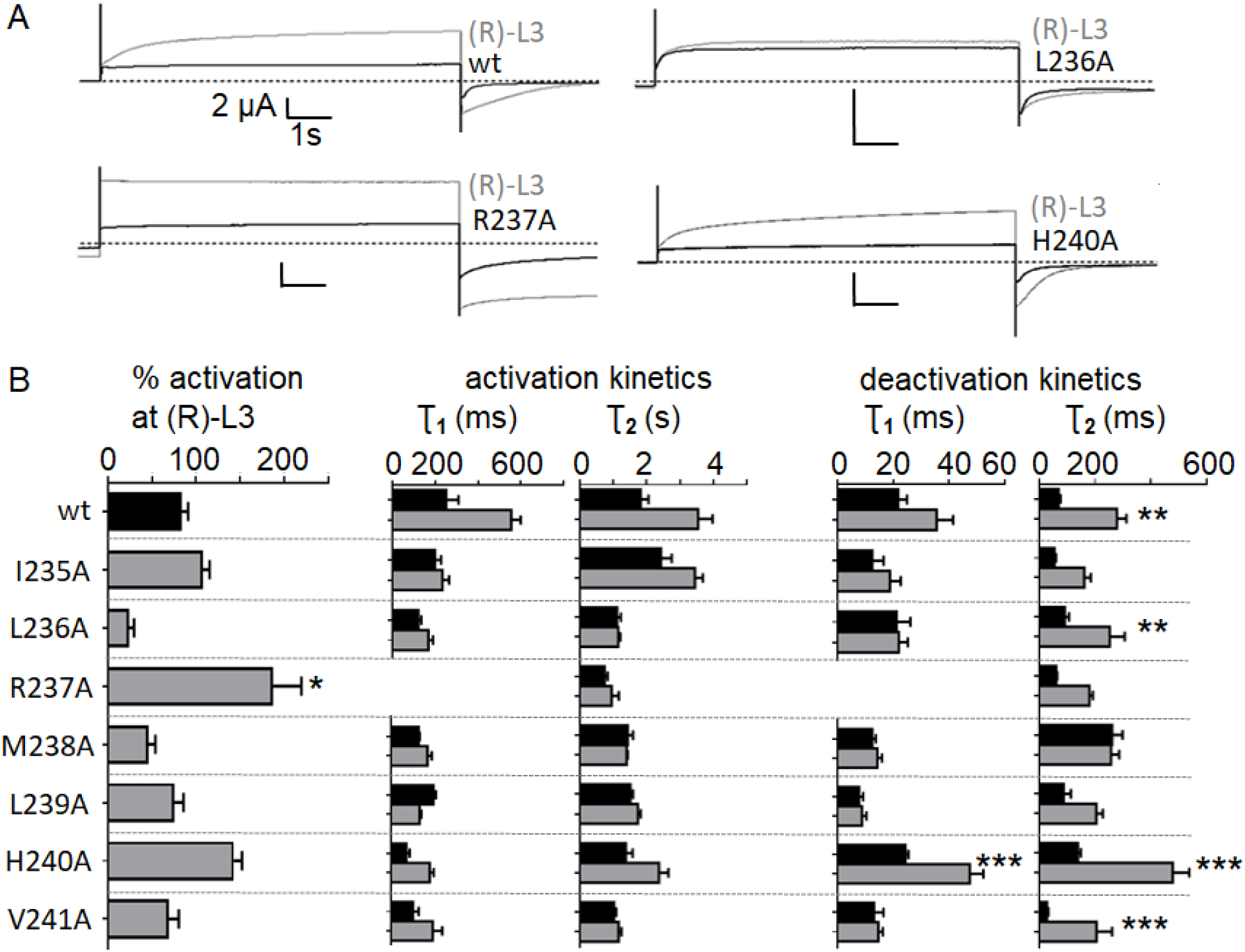
Sensitivity of S4 residues to (*R*)-L3 modulation. (A) Representative current traces of wildtype and mutant channels L236A, which was less sensitive to (*R*)-L3 and mutant channels R237A und H240A, which were more sensitive to the compound (n=13 to 18 ± SEM). **p<0.01, ***p< 0.001. (B) Current amplitude activation of K_v_7.1 channel mutants by 1 μM (*R*)-L3 expressed as percent change in current measured at the end of a 7-second pulse to +40 mV. Effect of (*R*)-L3 on K_v_7.1 channel mutant activation and deactivation rates. Kinetics was evaluated at +40 mV in both absence and presence of 1 μM (*R*)-L3 and fitted by two exponential function. Time constants for fast and for slow component of channel activation. Time constants for fast and for slow component of channel deactivation (n=13 to 18 ± SEM). *p<0.05, **p<0.01, ***p< 0.001.

In order to assess the relevance of key residue M238 in normal channel gating we conducted photo-crosslinking experiments. Modeling suggests a possible physical interaction of M238 with outer pore of a neighboring subunit by the S5 residue I271 in the activated state. This putative interaction was evaluated by incorporating the photoactivatable non-canonical amino acid p-azido-phenylalanine (AzF) at position Kv7.1-M238 (M238AzF) using the amber suppression method in HEK cells (Figure 5) ^34^. Application of high K^+^ buffer (137 mM) depolarized cells favoring voltage sensors in the activated up-state, which led to M238AzF-mediated inter-subunit photo-crosslinking, as shown by the formation of Kv7.1 multimers in Western blots (Figure 5E). These observations are consistent with the proposed interaction between M238AzF and a neighboring subunit pore domain (S5-S6), which is expected to result in covalent inter-subunit bonds. Further, as high K^+^ lead to depolarization of cells and subsequent channel inactivation, the M238AzF-I271 interaction may occur in open/inactivated channels. As an alternative but much less site-specific approach, we assessed the global incorporation of photo-methionine. Similar to our findings for M238AzF, photo-methionine incorporation allowed for photo-crosslinking and stable formation of K_v_7.1 multimers in Western blots in high K+ solution (Figure 5E). In summary, these two crosslinks indicate a M238-inter-subunit interaction under depolarizing conditions. This result is also consistent with a previous finding that M238 interacts with I271 of the neighboring subunit, that is involved in the coupling between the VSD and the pore ^35, 36^.

**Figure 5:**
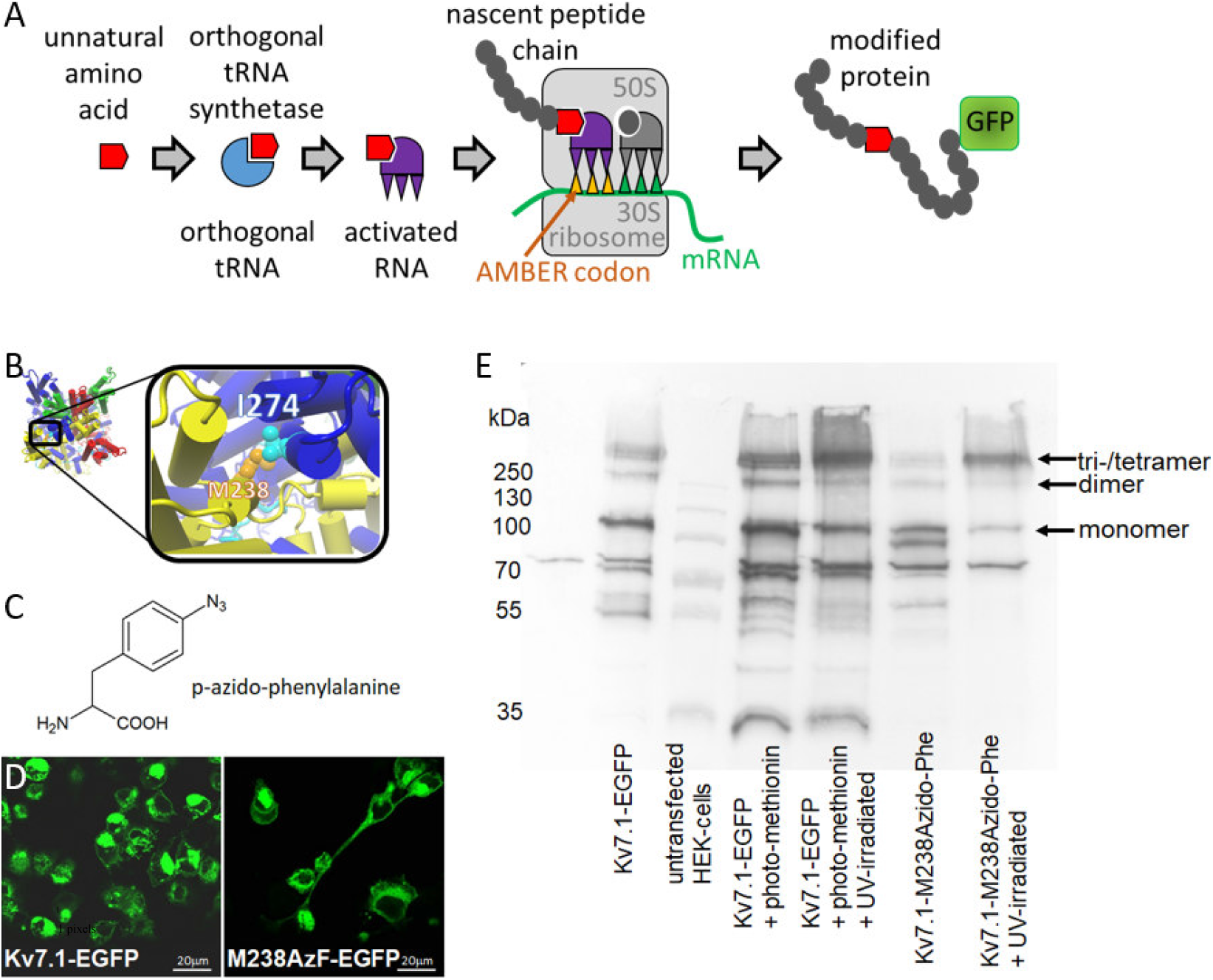
Photo-crosslinking of the photo-activatable non-canonical amino acid AzF leads to formation of multimers. (A) Using the amber suppression methodology for incorporation of non-canonical amino acids (ncAAs), we introduced the photo-activatable non-canonical amino acid AzF at M238 position into mutant Kv7.1-EGFP-M238AMBER-Stop. HEK cells were co-transfected with cDNA encoding Kv7.1-EGFP-M238amber-Stop, suppressor tRNA and AzF-tRNA synthetase. Addition of the ncAA amino acid p-acido-phenylalanine (AzF) at 0.5 mM to the cell culture medium allowed for incorporation of the ncAA. (B) I274 interacts with M238 from the adjacent subunit in activated state *in silico*. (C) Structure of photoactivatable ncAA AzF. (D) Successful incorporation and full-length protein expression was assayed by confocal EGFP imaging. (E) Kv7.1-EGFP-M238AzF expressing HEK cells were incubated in high K+ solutions (137 mM KCl,) leading to channels preferentially in depolarized (calculated V_m_ of about −8.9 mV) states. UV irradiation caused formation of multimers under high K+ / preferentially depolarizing conditions conditions consistent with the predicted interaction.

To connect the experimental data with structural analyses of the ligand/receptor complex and the protein movement, we used previously described models of different ion channel states for manual docking of (*R*)-L3 and subsequent molecular dynamics (MD) simulations ^37^. The used ion channel models are homotetrameric structures composed of four α-subunits (Mol A – Mol D) and display the receptor with an activated VSD and an open pore (AO), an activated VSD and a closed pore (AC) and a resting VSD with closed pore (RC). Based on the results from mutational analysis we built complexes of (*R*)-L3 and the three different receptor states (AO, AC, RC) by manually docking the ligand into the pocket between S4 and the S4S5 linker of Mol B with the indol-3-yl moiety orientated between the α-helices S5 and S6 from Mol A (Figure 6A-6C). The generated ligand/receptor complexes were energy minimized and embedded into lipid membranes surrounded by water molecules (Figure 6D). After a second annealing and energy-minimization step, MD simulations with a duration of 30 ns were performed 3 (RC, AC) or 5 (AO) times for the generated ligand/receptor complexes.

**Figure 6:**
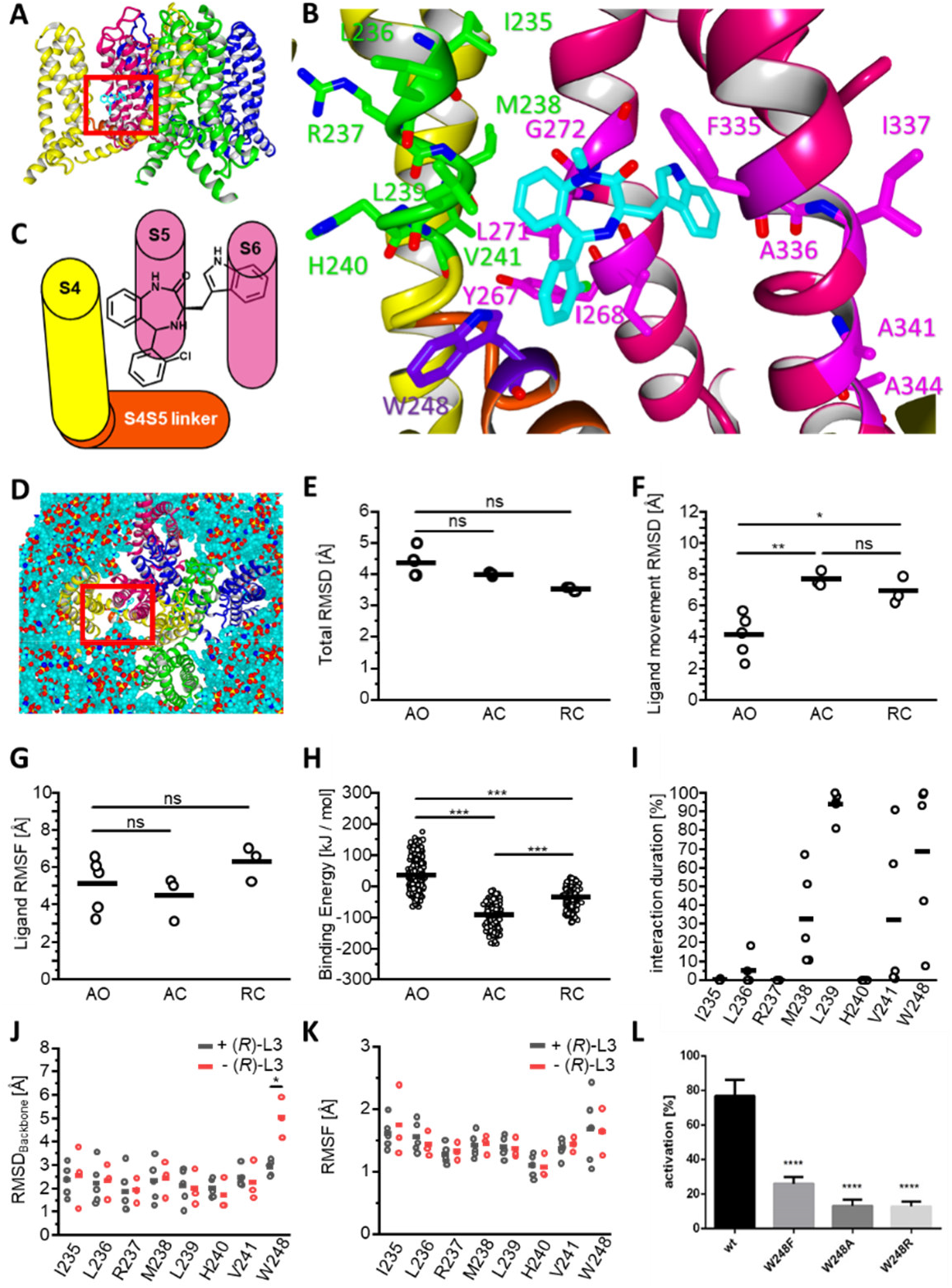
(A) Depiction of tetrameric KCNQ1 AO model with subunits colored in magenta (Mol A), yellow (Mol B), green (Mol C) and blue (Mol D). Proposed binding site of (*R*)-L3 is marked by red square. (B) Close-up depiction of (*R*)-L3 (CPK color coded with cyan for carbon) binding site with S4 (yellow) and S4S5 linker (orange) from subunit Mol B and S5 and S6 helix (magenta) from subunit Mol A. Previous analyzed amino acids from S5 and S6 with impact on (*R*)-L3 activity are colored in purple, while newly analyzed residues from S4 and S4S5 linker are colored in green or violet. (C) Schematic illustration of (*R*)-L3 binding site between S4 (yellow) and S4S5 linker (orange) from subunit Mol B and S5 and S6 (pink) from subunit Mol A. (D) Depiction of KCNQ1 AO model embedded in membrane for MD simulation. Subunits Mol A-D are colored like in 5A and (*R*)-L3 binding site is marked by red square. (E) Root mean square deviation (RMSD) of 30 ns MD simulations for complete modelled structures AO, AC and RC with (*R*)-L3 for every simulation. Means are shown as bar and have no significant differences indicated by ns. (F) RMSD of Ligand movement of (*R*)-L3 from start to end of the 30 ns MD simulation of KCNQ states AO, AC and RC. (G) (*R*)-L3 Root mean square fluctuation (RMSF) for 30 ns MD simulation with KCNQ1 state models AO, AC and RC. Mean differences are not significant (ns). (H) Binding Energy [kJ/mol] of (*R*)-L3 calculated for all simulation snapshots from 10 – 30 ns of MD simulations for KCNQ1 models AO, AC and RC. (I) Percentage duration of hydrophobic interaction between (*R*)-L3 and mutated amino acids from S4 and S4S5 linker over total MD simulation time of 30 ns. (J-K) Backbone RMSD and RMSF of mutated residues in absence / presence of (*R*)-L3. (L) activity of 1 μM (*R*)-L3 at wildtype KCNQ1 compared to mutations W248F, W248A and W248R.

For analysis of MD simulations, we used parameters like the root mean square deviation (RMSD), root mean square fluctuation (RMSF) and the movement correlation of residues expressed by dynamic cross correlation matrices (DCCMs). The RMSD can be calculated for multiple or single amino acids or even atoms and displays the total movement from the beginning of the simulation to the end, while RMSF displays the fluctuation around a mean position. After 30 ns of simulation, the total RMSD of the ligand/receptor complexes was constantly fluctuating around a steady state value indicating a stable formation. Although the total RMSDs decrease from the AO-complexes to the RC-complexes, the means were not significantly different (Figure 6E). On the contrary, the RMSD for (*R*)-L3, which displays the movement for the ligand away from the energy minimized starting position, was significantly reduced for the AO-complexes (Figure 6F). Since the RMSF of the ligand did not significantly differ, the different RMSDs must be caused by a movement of the complete ligand instead of a reorientation of ligand site chains (Figure 6G). All simulated complexes show a constantly fluctuating ligand RMSD after an equilibration time of 10 ns. Therefore, the binding energy for every ligand/receptor complex was calculated from 10 ns to 30 ns every 0.25 ns. The highest binding energies were observed in the AO-complexes indicating a better binding of (*R*)-L3 to this ion channel state (Figure 6H). Together with the reduced ligand RMSD in the AO state, these result lead to the conclusion of a favored binding to the fully activated AO receptor state.

Hence, we focused on the AO-ligand/receptor complex and analyzed interactions between (*R*)-L3 and the mutated amino acids of the lower S4 segment. Only hydrophobic interactions with L236, M238, L239 and V241 were detected. Additionally, a strong aromatic interaction between W248 from the S4S5 linker (Mol B) and the 2-chlorophenyl moiety was observed. Figure 6I shows the relative interaction duration over the complete simulation time, which indicates a strong influence of W248 on the binding. Consequently, we included W248 into further analyses. While distinct interactions with the receptor are needed for compound binding, the mechanism of activation is conducted by altered movements of the protein in presence of the ligand. To detect these movements, we performed three additional 30 ns MD simulations of the AO state model without the ligand for comparison. Surprisingly, the examined amino acids in S4 (Mol B) did not show significantly different RMSD or RMSF values indicating a similar behavior for the presence and absence of (*R*)-L3 (Figure 6J, 6K). In contrast, the RMSD of W248 from the S4S5 linker (Mol B) was significantly reduced without a significant change in the RMSF suggesting an inhibited movement of the amino acid in presence of (*R*)-L3.

Due to the frequently detected aromatic interactions with the ligand *in silico* and the altered behavior of W248 in the presence of (*R*)-L3 additional mutants W248F, W248A and W248R were generated. TEVC measurements revealed a dramatic loss of (*R*)-L3 activity for all mutants compared to wildtype emphasizing the involvement of W248 in the molecular mechanism of action (Figure 6L). Comparing the activity of (*R*)-L3 on aromatic mutant W248F with activity on W248A and W248R lead to the conclusion that not only the size but also the aromaticity of the amino acid at this position seems important for the agonistic mechanism.

While only minor differences are found on the scale of single interacting amino acids, major differences were observed in the communicational behavior of the binding site forming segments S4 (Mol B), S4S5 linker (Mol B), S5 (Mol A) and S6 (Mol A) (Figure 7A). To visualize the altered communication between the segments, DCCMs were calculated for all simulations and averaged depending on the presence or absence of (*R*)-L3 (Figure 7B – 7J). The dynamic cross correlation was calculated for a pair of Cα atoms over the complete MD simulation time and displayed movement behavior of the selected atoms in a range from −1 (fully anticorrelated movements) over 0 (no movement correlation) up to 1 (fully correlated movements). Although the S4 helix (Mol B) and the S4S5 linker (Mol B) are in close distance and on the same subunit their movements were not coordinated in the simulations without (*R*)-L3 (Figure 7B). In presence of the ligand a clear increase of coordinated movements could be observed between the lower S4 segment with the adjacent residues from the S4S5 linker (Figure 7C, 7D). On the contrary, the natural communication and the movement correlation between S4 (Mol B) and the corresponding S5 helix (Mol A) was nearly abolished (Figure 7E-7G). In accordance with the reduced correlation between S4 and S5 also the simultaneous movements of S5 and the S4S5 linker were reduced (Supplementary Figure 1A-1C). The presence of (*R*)-L3 generated also weak coupling between S4 (Mol B) and S6 (Mol A), while under normal condition no long-range coupling could be detected between these segments (Figure 7H-7J). In particular, the movement correlation between M238 and surrounding residues from S4 with the residues A336 – F340, which are directly interacting with (*R*)-L3, was increased. However, these differences were less pronounced than the coupling changes between S4, S4S5 linker and S5. A more pronounced elevation of coupling was achieved by (*R*)-L3 for the lower S6 helix with the whole S5 helix (SI Figure 1D-1F), while at the same time S6 was uncoupled from the movements of the S4S5 linker (SI Figure 1G-1I). Summarizing the movement coupling, the presence of (*R*)-L3 separated the movements of the VSD from the pore domain by decreasing the correlation between S4 and S5 and simultaneously increasing the correlation between S4 and S4S5 linker and between the inner pore domain forming helices S5 and S6.

**Figure 7:**
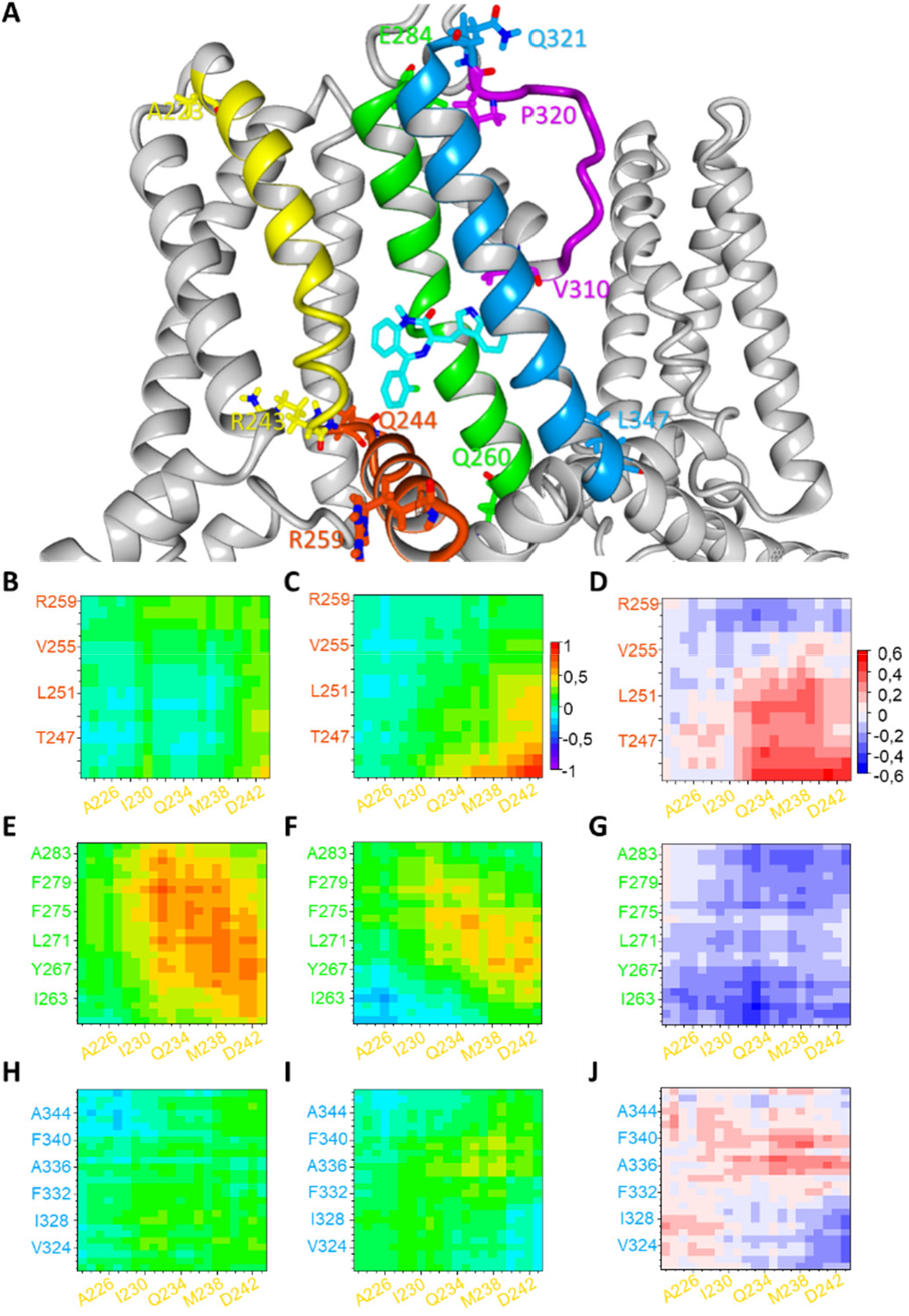
(A) (*R*)-L3 (CPK-color code with cyan for carbon) binding site between subunit Mol A and B with interacting secondary structures S4 (Mol B, yellow), S4S5 linker (Mol B, orange), S5 (Mol A, green), S6 (Mol A, blue) and selectivity filter (Mol A, purple). First and last amino acid of all secondary structures are shown in the same color. (B, C) Dynamic cross correlation matrix (DCCM) for S4 (Mol B, yellow) and S4S5 linker (Mol B, orange) in absence (B) and presence (C) of (*R*)-L3 from −1 (fully anticorrelated) over 0 (not correlated) to 1 (fully correlated). (D) Increase (positive values, red) and decrease (negative values, blue) of correlation between S4 and S4S5 linker residues depending on presence of (*R*)-L3. (E, F) DCCM for S4 (Mol B, yellow) and S5 (Mol A, green) in absence (E) / presence (F) of (*R*)-L3. (G) Increase (positive values, red) and decrease (negative values, blue) of correlation between S4 and S5 residues depending on presence of (*R*)-L3. (H, I) DCCM for S4 (Mol B, yellow) and S6 (Mol A, blue) in absence (H) / presence (I) of (*R*)-L3. (J) Increase (positive values, red) and decrease (negative values, blue) of correlation between S4 and S6 residues depending on presence of (*R*)-L3.

Since (*R*)-L3 increases K_v_7.1 current amplitude, an allosteric coupling between the binding site forming structures and the adjacent selectivity filter residues can be assumed (Figure 8A). Hence, RMSDs of amino acids V310 – P320 (Mol A) and the coupling of these with S4, S4S5 linker, S5 and S6 were analyzed (Figure 8B-8O). Comparing the residue backbone RMSD in absence and presence of (*R*)-L3, the amino acids V310 – T312 and K318 – P320 showed a moderate, but not significant increased movement when the ligand is bound (Figure 8B). However, the averaged backbone RMSD for V310 – P320 was significantly increased suggesting an elevated movement of the whole selectivity filter segment (Figure 8C). Similar to previous results for S4, the movement correlation of S4 with the selectivity filter decreased. In particular, the correlation between M238 and the lower residues was extinguished by the ligand (Figure 8D-8F). Contrary to this, a slightly increased movement correlation occurred between the S4S5 linker and the lower amino acids of the selectivity filter (Figure 8G-8I). While the outer structures showed only minor correlation differences with V310 – P320, the two inner helices S5 and S6 nearly swap their behavior towards the selectivity filter (Figure 8J-8O). S5 loosed correlation especially to the amino acids T312 – K318, while S6 achieves a higher grade of correlation with the lower selectivity filter. Especially F339, which is directly connected to the indol-3-yl moiety of (*R*)-L3 via aromatic interactions, amplified its movement coupling with V310 – I313. In summary, (*R*)-L3 raised the mobility of V310 – P320 and altered the coupling of these residues by enhancing the movement correlation with S6 and simultaneously decreasing it with S4 and S5.

**Figure 8:**
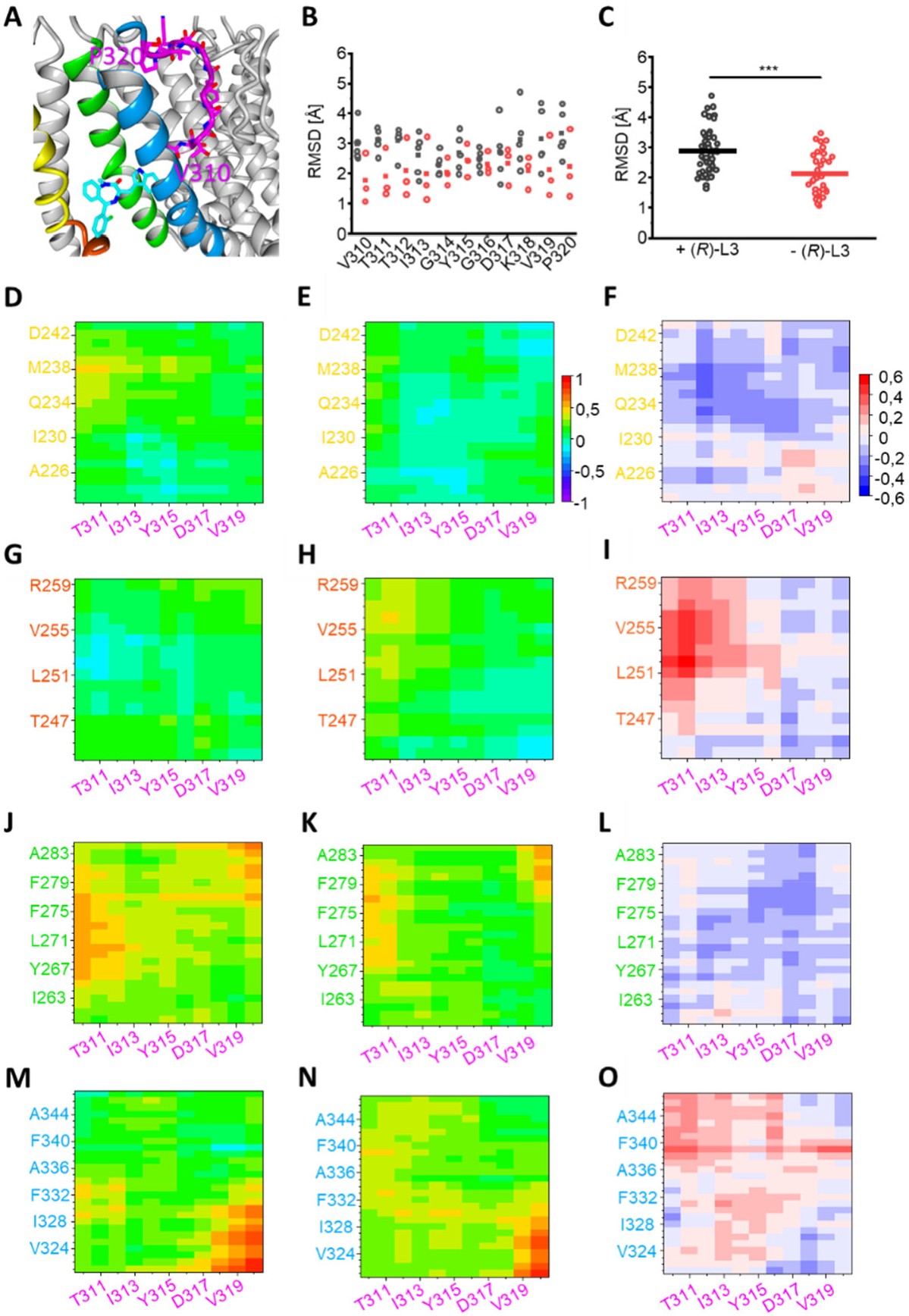
(A) Close-up depiction of (*R*)-L3 binding site with S4 (yellow), S4S5 linker (orange), S5 (green), S6 (blue) and selectivity filter (purple) with all amino acids from V310 to P320. (B, C) RMSD of selectivity Filter residues V310 – P320 individually (B) and in total (C) in absence (black) and presence (red) of (*R*)-L3. Means are given as squares (B) or bars (C). (D, E) Dynamic cross correlation matrix (DCCM) for S4 (Mol B, yellow) and selectivity filter (Mol A, purple) in absence (D) and presence (E) of (R)-L3 from −1 (fully anticorrelated) over 0 (not correlated) to 1 (fully correlated). (F) Increase (positive values, red) and decrease (negative values, blue) of correlation between S4 and selectivity filter residues depending on presence of (*R*)-L3. (G, H) DCCM for S4S5 linker (Mol B, orange) and selectivity filter (Mol A, purple) in absence (G) / presence (H) of (*R*)-L3. (I) Increase (positive values, red) and decrease (negative values, blue) of correlation between S4 and selectivity filter residues depending on presence of (*R*)-L3. (J, K) DCCM for S5 (Mol A, green) and selectivity filter (Mol A, purple) in absence (J) / presence (K) of (*R*)-L3. (L) Increase (positive values, red) and decrease (negative values, blue) of correlation between S4 and selectivity filter residues depending on presence of (*R*)-L3. (M, N) DCCM for S6 (Mol A, blue) and selectivity filter (Mol A, purple) in absence (M) / presence (N) of (*R*)-L3. (O) Increase (positive values, red) and decrease (negative values, blue) of correlation between S4 and selectivity filter residues depending on presence of (*R*)-L3.

## Discussion

Since K_v_7.1 channels are key channels in several physiological processes, functional impairment by mutations or by drug side effects can lead to severe diseases including arrhythmias, diabetes and others ^4, 10, 38–47^. Thus, pharmacological modulation of K_v_7.1 channels holds the potential for specific drug development. In the present study, we analyzed the binding site and mechanism of action of the known K_v_7.1 modulator (*R*)-L3, that slows down channel activation and deactivation and increases current amplitude ^48^ (Figure 1). Current amplitude, inactivation, activation and deactivation kinetics are dependent on the permeating ion, via allosteric effects within the pore domain ^18, 27^. Using this phenomenon, we exchange the permeating ion to discriminate between kinetic (*R*)-L3 effects and current amplitude (*R*)-L3 effects (within the pore). Indeed, the increment of current amplitude by the modulator is clearly dependent on the permeating ion (Figure 2). Residues in the S5-S6 pore domain have been identified to be involved in both, formation of the (*R*)-L3 binding site ^23^ as well as formation of an allosterically linked cluster among lower selectivity filter. Thus, S5 and S6 determining current amplitudes and inactivation ^18, 27, 49^. Therefore, the modulator may allosterically influence the conduction pathway to allow for increased conduction rates. In contrast, (*R*)-L3 effects on current kinetics are similar in high Na^+^, K^+^ and Rb^+^ (Figure 2H,I) which indicates that the activity of the modulator can be functionally separated in a stimulation effect of current amplitude via modulation of the conduction pathway and a second (*R*)-L3 effect on current kinetics by a different mechanism.

The (*R*)-L3 impact on current kinetics could result from altered voltage sensor movement. To confirm this hypothesis, voltage clamp fluorometry (VCF) was performed and showed that the (*R*)-L3 effects are followed by parallel left shifted GV and FV curves (Figure 3). This finding is consistent with mathematical modeling suggesting that the modulator favors opened over closed states ^23^. For the coupling of VSD movements to the pore domain PIP_2_ was recently described as important modulator ^50^. Interestingly, voltage dependent CiVSP-depletion of PIP_2_ does not reverse the effect of the compound on the FV indicating that (*R*)-L3 does not require PIP_2_ to modulate K_v_7.1 gating (Figure 3C). Thus, (*R*)-L3 seems to exert a direct impact on the voltage-sensing domain.

If this hypothesis is correct, then binding of (*R*)-L3 between the pore domain and the VSD would be indicated. By inspection of the K_v_7.1 homology model the identified S5-S6 residues suggest a binding site from the interface of S5 and S6 (indol-3-yl sidechain) to the outer S5 (benzodiazepinic part). The outer S5 of K_v_ channel subunit is expected to be located in close proximity of the primary voltage sensing helix S4 of the adjacent subunit, suggesting that (*R*)-L3 may interact with S4 as well. Usage of classical mutagenesis scanning of the lower S4 identifies several residues modulating the activation of K_v_7.1 channels. The most dramatic effect was seen with mutation at M238 whose mutation significantly blunted (*R*)-L3 effects (Figure 4A, 4B). Other mutants impaired compound effects on either activation (L236A, R237A, L239A, V241A) or on deactivation (I235A, L239A) or on current amplitudes (L236A, R237A, H240A) but not on both parameters like M238A. Therefore, M238 may be considered a key residue for (*R*)-L3 activity.

Recently, a two-stage hand-and-elbow gating mechanism for K_v_7.1 was proposed. S4 movement towards a fully-activated conductive state depends on an elbow-like hinge between S4 and S4S5 linker that engages with the pore of the neighboring subunit to activate conductance ^35^. Based on this open state model interaction of M238 with the neighboring S5-S6-domain can be predicted *in silico*. In order to test this hypothesis *in vitro* we adapted a photo crosslinking approach and show crosslinking of M238AzF with the neighboring subunit in open-state (Figure 5). Thus, M238 structurally interacts in the open state with the outer pore domain and may potentially modulate its function.

In order to evaluate the binding site further we used different K_v_7.1 channel models (AO, AC and RC state) and manually docked the (*R*)-L3 molecule assuming close proximity of key residues. Thus guided by the experimentally identified key residues in S5/S6 ^23^ and S4 (M238) we developed an *in silico* model (Figure 6). In MD simulations (*R*)-L3 was stably coordinated over 24 ± 5.45 nanoseconds (n = 5) only to the AO model to the postulated binding pocket in close proximity of the key residues suggesting favorable binding only in the fully activated state. By inspection of the model complex, residue W248 in the S4S5 linker was identified as a likely interaction partner. In order to verify our model, we mutated the natural tryptophan and found that it is highly important for the activity of the modulator on the K_v_7.1 channel (Figure 6D). This finding largely strengthens the model and suggests its validity. Therefore, extensive evidence suggests that (*R*)-L3 binds in a binding pocket formed by S4, S4-S5 linker and the pore domain S5/S6 of the neighboring K_v_7.1 channel subunit. This is exactly the key region of electro-mechanical coupling in K_v_7.1-voltage-dependent activation described very recently ^35^. (*R*)-L3-uncoupling of key residue M238 from the neighboring S5-S6-domain and the lower K^+^-coordinating residue T331 in the lower selectivity filter in the open state may allow for higher conduction upon (*R*)-L3. Stabilization of interactions between the hand-like C-terminus of the VSD-pore linker (S4-S5L) with the pore (S5-S6) stabilizes the fully-activated state conformation as suggested by kinetic Markov modelling ^48^. In addition, this hypothesis is in good accordance with the correlated movement of M238 and the segments S4S5 linker and selectivity filter. Without (*R*)-L3 the movement of M238 correlates with residues from the selectivity filter (Figure 8D; residues V310-Y315), especially with T312, while no coordinated movement can be seen with the S4-S5 linker (Figure 7C). In presence of (*R*)-L3 this behavior towards the linker and the selectivity filter is reversed underlining the separation of VSD and pore domain by the compound.

Summarizing, this study provides evidence for two individual molecular mechanisms allowing K_v_7.1 channel activation by (*R*)-L3: Firstly, current amplitudes are increased due to compound binding to the pore domain to allosterically modulate the ion pathway. Secondly, K_v_7.1 channels may be trapped in open states by stabilization of the voltage sensors in the activated state (up-state).

## Materials and Methods

### Mutagenesis and heterologous expression of potassium channels

Site-directed mutagenesis of K_v_7.1 cDNA (GenBank Acc. No. NM_000218) subcloned into the pXOOM vector was performed using standard mutated oligo extension PCR. Mutations were verified by automated DNA sequencing. The constructs were linearized with NheI, and *in vitro* transcriptions were performed using the mMessage mMachine kit (Ambion) with T7 polymerase according to manufacturer’s instructions. Quality of synthesized RNA was checked by gel electrophoresis and quantified by spectrophotometry.

Oocytes were isolated from ovarian lobes of *Xenopus laevis* and enzymatically digested with collagenase (Type II, Worthington, 1 mg/ml in calcium-free Barth’s solution) for about 1.5-2 h, as previously described. Stage IV or V oocytes were injected with 5 ng cRNA encoding either wildtype or mutant K_v_7.1 subunits and stored for 4 days at 18 °C in Barth’s solution containing (in mmol/L): 88 NaCl, 1.1 KCl, 2.4 NaHCO_3_, 0.3 Ca(NO_3_)_2_, 0.3 CaCl_2_, 0.8 MgSO_4_, 15 HEPES-NaOH, penicillin-G (31 mg/L), gentamycin (50 mg/L), streptomycin sulfate (20 mg/L), pH 7.6 before electrophysiological recording.

### Electrophysiology

Whole-cell currents in oocytes were recorded at room temperature by two-electrode voltage clamp (TEVC) using a Turbo Tec 10CD amplifier (NPI Electronic GmbH, Tamm, Germany) and an ITC-16 interface combined with GePulse software (Michael Pusch, Genova, Italy). Recording pipettes pulled from borosilicate glass were filled with 3 M KCl and had resistances of 0.4-1 MΩ. Recordings were performed in ND96 solution containing (in mM) 96 NaCl, 4 KCl, 1 CaCl2, 1.8 MgCl2, 5 HEPES, pH 7.2-7.4 with 0.1 % dimethylsulfoxide. For experiments with high extracellular potassium or rubidium concentrations, standard ND96 solution was replaced by 100 mM KCl and 100 mM RbCl solution, respectively. (*R*)-L3 was added to each bath solution to give a final concentration of 1 μM, from a 10 mM stock solution in DMSO. To determine the effects of (*R*)-L3 on wildtype and mutant K_v_7.1 channels, currents were recorded by repetitive 7-second pulses to potentials of −100 mV to +60 mV, applied in 20 mV increments and then held for a 3-second tail pulse at −120 mV before returning to the holding potential of −80 mV.

### Data Analysis

Data were analyzed with the custom program Ana and GraphPad Prism 5.01 software (GraphPad Software, San Diego, California). All experimental results are presented as mean ± SEM with n specifying the number of independent experiments. Statistical analysis was performed with student’s t-test (unpaired) or ANOVA. The activation curves for wildtype and mutant K_v_7.1 channels were determined from tail currents at hyperpolarized potentials by plotting the normalized peak tail current amplitudes versus the respective test potential. The resulting curves were fitted to a Boltzmann equation that gave an adequate fit with the two parameters V_1/2_ (voltage of half maximal activation) and the slope factor k. Activation kinetics was obtained by double-exponential fits to the currents yielding fast and slow time constants of activation. Deactivation rate constants were also obtained by double exponential fits to the tail currents in the absence and presence of (*R*)-L3 and plotted against the applied membrane potential.

### Methods of Voltage Clamp Fluorometry (VCF)

Oocytes were injected with 9.2 ng of cRNA encoding of psWT K_v_7.1 (C214A/G219C/C331A) alone or simultaneously with 2.3 ng of cRNA encoding wt CiVSP. After incubation for 3-7 days at 18 C, the cells were labeled in high K^+^ labeling solution (98 mM KCl, 1.8 mM CaCl_2_, 5 mM HEPES, pH 7.6) with 10 μM Alexa 488 C_5_-maleimide (Molecular Probes) for 45 minutes on ice, washed with ND96 solution (96 mM NaCl, 4 mM KCl, 1.8 mM CaCl_2_, 1 mM MgCl_2_, 5 mM HEPES), and kept on ice until recording to minimize recycling of labeled channels. VCF recordings were carried out in ND96 solutions +/− (*R*)-L3. Fluorescence recordings were performed using a Leica DMLFS upright fluorescence microscope with a FITC filter cube. Emission from the animal pole was focused onto a pin 20A photodiode (OSI Optoelectronics) and amplified using an EPC10 (HEKA) patch amplifier and analog filtered at 200 Hz. Simultaneous two electrode voltage clamp recordings were measured using a Dagan CA1-B amplifier in TEVC mode. Current and fluorescence signals were digitized at 1 kHz.

### Cell culture, transfection, photo-crosslinking and cell lysis

HEK293T cells were cultured in DMEM (Sigma-Aldrich) supplemented with 2 mM L-glutamine (Sigma-Aldrich), 100 U/ml penicillin (Sigma-Aldrich), 10% fetal bovine serum (Biochrom) and 100 μg/ml streptomycin (Sigma-Aldrich) at 37°C / 5% CO_2_. For transfection, cells were grown to 60-70% confluency in a 3.5 cm dish. For amber suppression experiments, the plasmid DNA ratio used for transfection was 1/1/0.2 (K_v_7.1/suppressor tRNA/AzF aaRS). Transfection was performed using FuGene HD (Promega) according to the manufacturer’s protocol. For the double transfection 625 ng K_v_7.1 DNA, 625 ng suppressor tRNA DNA and 125 ng aaRS DNA were used per 3.5 cm dish. For transfection of wildtype (wt) K_v_7.1, 1 μg of DNA was used per 3.5 cm dish. Non-transfected cells served as controls. Cells transfected with K_v_7.1 amber mutants were grown in the presence of 0.5 mM AzF (Chem-Impex) in the medium. Cells expressing K_v_7.1-M238AzF were harvested by trypsinization 24-48 hours post transfection, centrifuged, exposed to 254 nm UV-light for 3 minutes and pelleted again. The pellet was solubilized for 30 minutes at 4°C and cell lysates were purified using the μMacs GFP isolation kit (Miltenyi Biotec) according to the manufacturer’s protocol. SDS-PAGE was performed and proteins were transferred to nitrocellulose membranes. Following protein transfer, membranes were stained with Ponceau S solution (Sigma-Aldrich) and destained with Tris Buffered Saline (TBS: 20 mM Tris, 150 mM NaCl). For detection of K_v_7.1, the membrane was immunoblotted with monoclonal K_v_7.1 antibody (sc-365186, Santa Cruz Biotechnology) 1:200 in TBS-T (TBS + 0.1 % Tween).

### Molecular Modeling

Molecular modelling was performed using YASARA 19 and OriginPro 2019 for data analysis. K_v_7.1 structures of channel states AO, AC and RC were adopt from Kuenze *et al*. ^37^. PDB files of different channel states are comprised of four identical KCNQ1 subunits named Mol A – D. Structure of (*R*)-L3 was constructed with YASARA 19 with defined stereochemistry and energy minimized using AMBER14 force field. Based on results from previous and currently performed mutagenesis studies, energy minimized (*R*)-L3 was manually docked between Mol A and Mol B in close proximity to all interacting amino acids from S4 (Mol B), S5 (Mol A) and S6 (Mol A) followed by a second energy minimization using AMBER14. For MD simulation in a lipid membrane, the provided YASARA 19 macro md_runmembrane.mcr was used and the following parameters were adapted: Membrane composition was set to 1/3 phosphatidyl-ethanolamine (PEA), 1/3 phosphatidyl-choline (PCH) and 1/3 phosphatidyl-serine (PSE). All polar head groups are 1-palmitoyl- and 2-oleoyl substituted. Simulation speed was set to fast (simulation time steps of 2*2.5 fs) and simulation time was set to 30 ns. Process of simulation was documented by screenshots every 0.25 ns leading to 120 screenshots in total for every simulation. RMSD, RMSF and DCCM were calculated with provided YASARA macros md_analyze.mcr and md_analyzeres.mcr. Full simulation and analysis were conducted using AMBER14 force field.

Where relevant, significance of mean differences for simulation data was tested by one-way-ANOVA and posthoc mean comparison Tukey test indicated by * for p < 0.05, ** for p < 0.01, *** for p < 0.001 and ns (not significant) for p > 0.05.

## Supporting information

Supplemental Informations

## Acknowledgements

We thank Geoffrey Abbott for providing K_v_7.1 mutant channel constructs. This work was supported by R01 HL126774 from NIH to JC and an AHA fellowship (11PRE5720009) to MZ.

## Author contributions

MM, NR, SB, EW, MZ and ZB performed experiments, JAS, GS performed 3D simulations, NSS, JC, BW, JAS and GS experimental design, MM, EW, NS, NSS, BW, ND, MD, JAS and GS wrote the manuscript. All Authors critically revised the manuscript.

