## Supplemental Informations for "An Activator Locks Kv7.1 Channels Open by Electro-Mechanical Uncoupling and Allosterically Modulates its Pore"

**Running title: RL3 Binding and Activation**


Melina Möller[1,*], Julian A. Schreiber[1,2,*], Mark Zaydman[3], Zachary Beller[3], Sebastian Becker[1], Nadine Ritter[1,4], Eva Wrobel[1], Nathalie Strutz-Seebohm[1], Niels Decher[5], Jianmin Cui[3], Nicole Schmitt[6], Martina Düfer[2,4], Bernhard Wünsch[2,4], and Guiscard Seebohm[1,4,π]

[1]*Institute for Genetics of Heart Diseases (IfGH), Department of Cardiovascular Medicine, University Hospital Münster, D-48149 Münster, Germany.*

[2]*Institute of Pharmaceutical and Medicinal Chemistry, University of Münster, Corrensstr. 48, Münster, D-48149, Germany.*

[3]*Department of Biomedical Engineering, Center for the Investigation of Membrane Excitability Disorders, Cardiac Bioelectricity and Arrhythmia Center, Washington University, St. Louis, MO 63130*

[4]*Chembion, University of Münster, D-48149 Münster, Germany.*

[5]*Institute of Physiology and Pathophysiology, Vegetative Physiology, Philipps-University of Marburg, Deutschhausstr. 1-2, 35037 Marburg, Germany.*

[6]*Department of Biomedical Sciences, University of Copenhagen, Copenhagen, Denmark.*

* These authors contributed equally.

π Corresponding author:
Prof. Dr. Guiscard Seebohm
Tel. +49 (0)251/83-58255
Fax +49 (0)251/83-58257


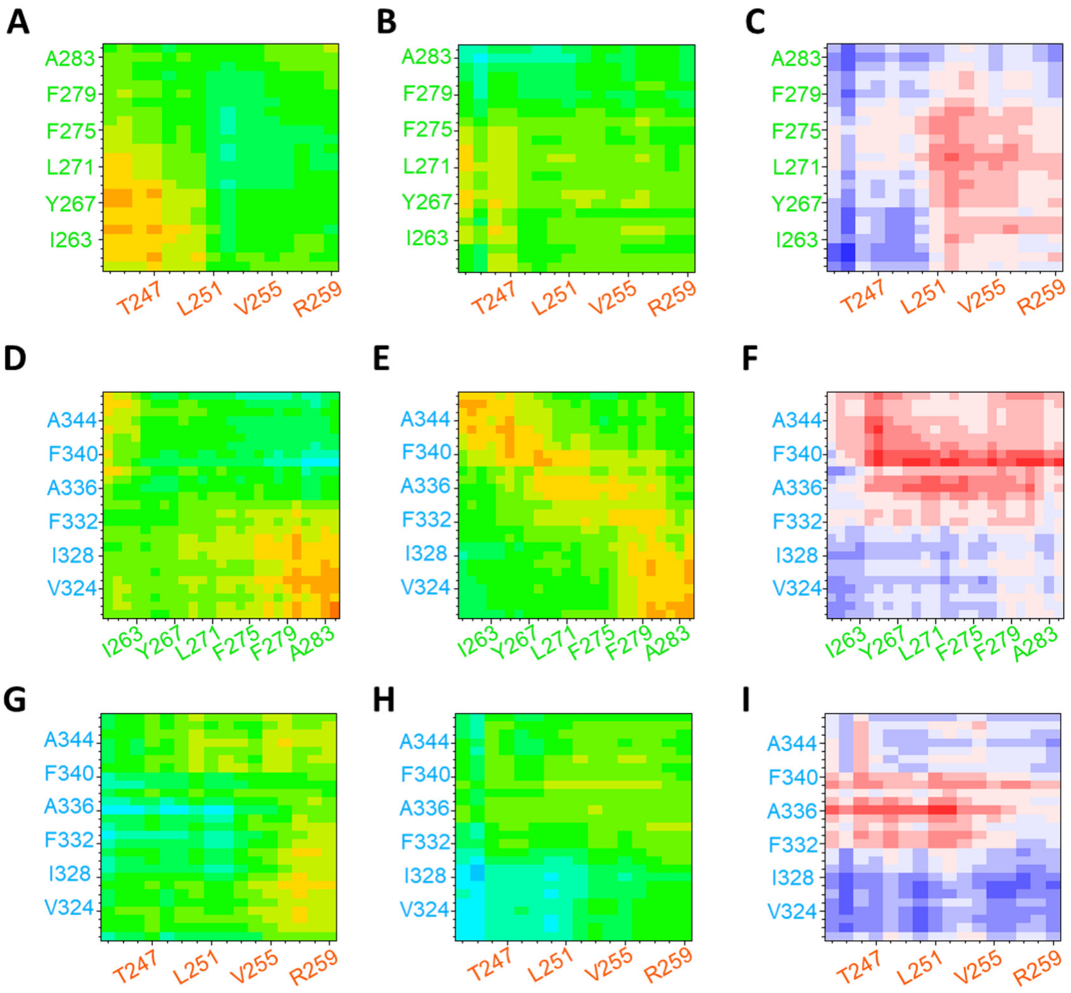


Supplementary Figure 1: (A, B) Dynamic cross correlation matrix (DCCM) for S5 (Mol A, green) and S4S5 linker (Mol B, orange) in absence (A) and presence (B) of (*R*)-L3 from -1 (fully anticorrelated) over 0 (not correlated) to 1 (fully correlated). (C) Increase (positive values, red) and decrease (negative values, blue) of correlation between S5 and S4S5 linker residues depending on presence of (*R*)-L3. (D, E) DCCM for S6 (Mol A, blue) and S5 (Mol A, green) in absence (D) / presence (E) of (*R*)-L3. (F) Increase (positive values, red) and decrease (negative values, blue) of correlation between S6 and S5 residues depending on presence of (*R*)-L3. (G, H) DCCM for S6 (Mol A, blue) and S4S5 linker (Mol B, orange) in absence (G) / presence (H) of (*R*)-L3. (I) Increase (positive values, red) and decrease (negative values, blue) of correlation between S6 and S4S5 linker residues depending on presence of (*R*)-L3.
